# Algorithmic and data modeling: Will algorithmic modeling improve predictions of traits evaluated on ordinal scales?

**DOI:** 10.1101/2020.10.07.329466

**Authors:** Zhanyou Xu, Andreomar Kurek, Steven B. Cannon, Williams D. Beavis

## Abstract

Selection of markers linked to alleles at quantitative trait loci (QTL) for tolerance to Iron Deficiency Chlorosis (IDC) has not been successful. Genomic selection has been advocated for continuous numeric traits such as yield and plant height. For ordinal data types such as IDC, genomic prediction models have not been systematically compared. The objectives of research reported in this manuscript were to evaluate the most commonly used genomic prediction method, ridge regression and it’s equivalent logistic ridge regression method, with algorithmic modeling methods including random forest, gradient boosting, support vector machine, K-nearest neighbors, Naïve Bayes, and artificial neural network using the usual comparator metric of prediction accuracy. In addition we compared the methods using metrics of greater importance for decisions about selecting and culling lines for use in variety development and genetic improvement projects. These metrics include specificity, sensitivity, precision, decision accuracy, and area under the receiver operating characteristic curve. We found that Support Vector Machine provided the best specificity for culling IDC susceptible lines, while Random Forest GP models provided the best combined set of decision metrics for retaining IDC tolerant and culling IDC susceptible lines.

## Introduction

Soybean iron deficiency chlorosis (IDC) is associated with yield losses of 340 million tons, worth an estimated $120 million per year in the upper Midwestern U.S. [1]. Breeding for IDC tolerance using markers linked to alleles at quantitative trait loci (QTL) has not been successful. The trait is evaluated on an ordinal scale (Fig 1), but is difficult to obtain repeatable scores due to ephemeral and environmental conditions required for its expression [2,10]. None-the-less, due to its economic impact, dozens of experiments have identified more than 100 QTLs including 41 IDC QTLs from bi-parental linkage mapping studies, 50 QTLs from connected network mapping studies, and 88 QTLs from GWAS studies SoyBase (https://soybase.org/) ([2-9]). In addition, 835 candidate genes in the IDC resistant line Clark (PI548553) were identified by transcriptome sequencing [14].

**Fig 1.**
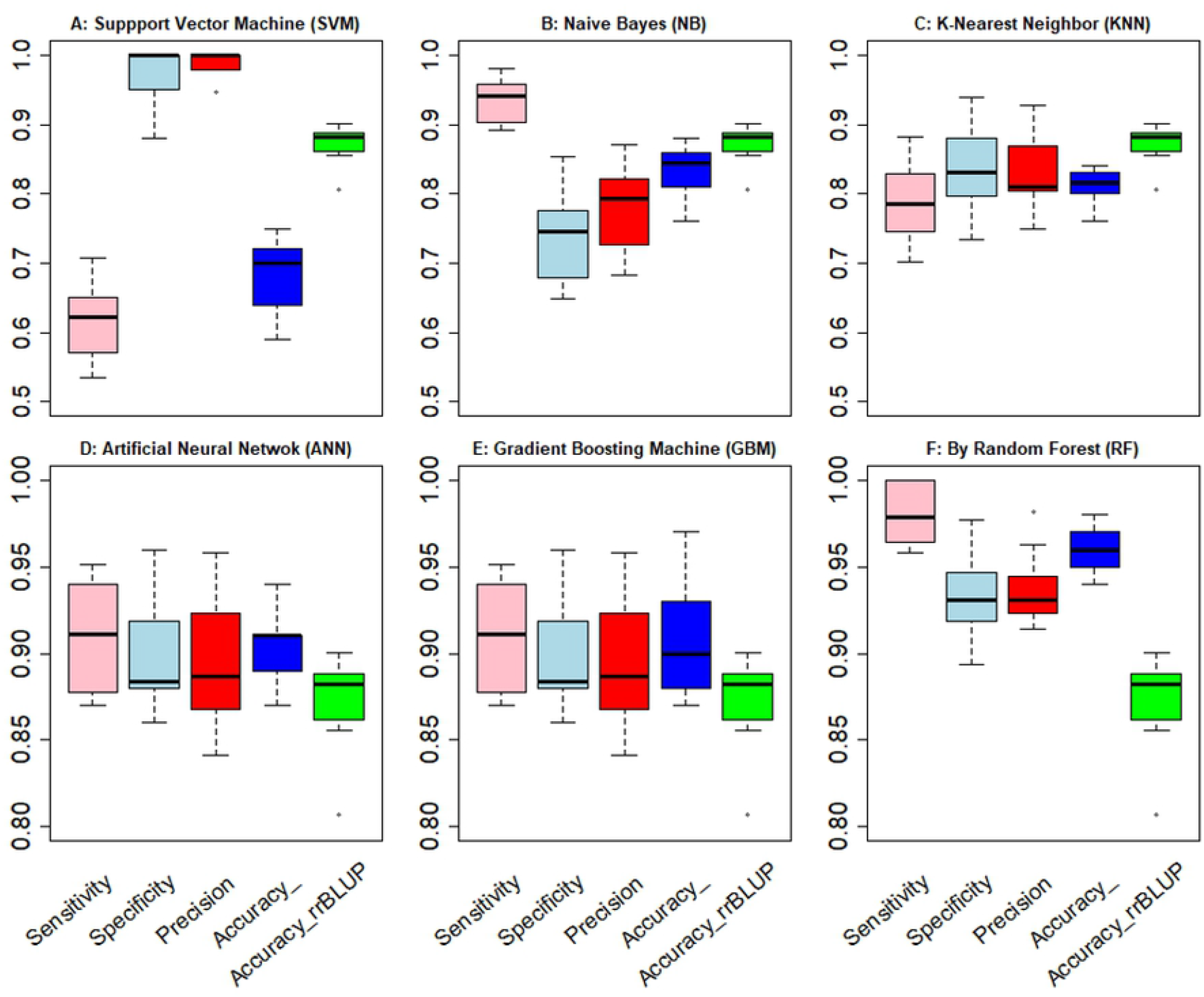
Boxplot of selection sensitivity, discard specificity, selection precision, and decision accuracy of six models. The four box plots in each model from left to right are the selection sensitivity, discard specificity, selection precision, and overall accuracy, the fifth box is from rrBLUP as the baseline for comparison. Y-axis are the accuracies from 0 to 1.

Since the first QTL studies for IDC, soybean breeders have been attempting to use marker alleles to stack desirable IDC QTL alleles without much success. In contrast to biologicial pest resistance, there have been few consistently identified QTLs with estimated large effects for IDC. For example, there is a consistently identified allele for resistance to soybean cyst nematode, derived from PI88788 that has been introgressed into 95% of the soybean commercial varieties [11]. The only ‘large effect’ IDC QTL was mapped in a population between Anoka x A7, which identified a QTL that explained more than 70% of the phenotype variation among the sample of progeny from that particular cross [12, 13]. However, no reports have found that this major IDC QTL provides resistance in other genetic backgrounds. Indeed, identified IDC QTL appear to be highly dependent on genetic background [4]. To summarize, tolerance to IDC is an important economic trait with ephemeral expression making it an ideal trait for application of MAS, but QTL based MAS for IDC tolerance has been ineffective because the genetics underlying the expression of the trait are affected by many genes and environmental factors [15, 16].

Genomic selection (GS) is practiced with various genomic prediction (GP) models that have been developed in conjunction with high-density genotyping and whole-genome sequencing technologies [17]. GP models use a training set of sampled genotypes (individuals, lines, hybrids, etc.) that have been both genotyped and phenotyped to train a model [17, 18]. Since GS was first proposed in 2001 results from applications to complex polygenic traits indicate that GS provides greater responses to selection than QTL based MAS in both empirical and simulation experiments [20, 21, 33, 34, 40, 41].

Most studies that compare GP models use prediction accuracy as a metric to assess relative performance. Prediction accuracies of GP models applied to *in vivo* experiments are estimated using the Pearson correlation coefficient, designated *r*, between genome estimated breeding values and the observed phenotypic values [19-29]. The expected prediction accuracy E (*r*_*mg*_) is defined as a function of sample size (N), heritability of the trait of interest (*h*^2^), and number of effective markers *M*_*e*_ associated with the trait of interest [30, 31]:

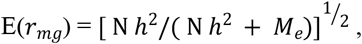

where h^2^ is the heritability of the trait of interest and is considered a fixed value for the trait, N is the sample size of evaluated genotypes and *M*_*e*_ refers to an idealized concept of independent chromosome fragments, with each linkage block containing net positive or negative effects from QTL-marker pairs [31, 32]. When the marker density is high enough to cover every functional gene in the genome, *M*_*e*_ can be estimated on the basis of the effective population size and the genome size [33, 34].

To estimate the prediction accuracy, four statistical assumptions are required [31]: 1) the marker effects can be derived from simple linear regression; 2) each marker–QTL pair is independent of other marker–QTL pairs; 3) different marker–QTL pairs have equal variances; and 4) each marker–QTL pair is in complete linkage disequilibrium (LD). For the currently used soybean high density SNP chip, not all markers are in LD with flanking QTLs. The equation was updated by Lian et al [32] to retain the first three assumptions but to relax the fourth assumption by adding an extra variable (*r*^2^_*MM*/2_) [32] :

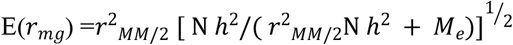

where *r*^2^_*MM*/2_ equals to the mean squared correlation between a marker and QTL when the QTL is assumed to be at the midpoint between two markers.

Prediction accuracies have been estimated from GP models applied to several crop species. For example, analysis of 1,536 SNPs and 647 barley lines for four traits evaluated over five years ranged from 0.03 to 0.99. Differences among the prediction accuracies indicate that higher trait heritability in the training population and simpler trait architectures were associated with greater estimated prediction accuracies that [35]. In maize, progeny of 969 bi-parental populations were genotyped with 31 to 119 (with a mean of 70) polymorphic SNPs. The estimated prediction accuracies ranged from -0.59 to 1.03 with a mean of 0.45 for grain yield [32]. The authors concluded that it is difficult to predict the accuracy, *r*_*MG*_, but ta rule of thumb based on *r*^2^_*MM*/2_(*Nh*^2^)^½^ >8 can help.

Two distinct modeling approaches have been used for GP. These approaches are distinct enough that they have been termed statistical “cultures” by Brieman [42]. The “data modeling” approach uses pre-defined, distribution-based models. The majority of developed GP models fall within this category. A best linear unbiased prediction algorithm for ridge-regression (rrBLUP) is the most widely used GP model, largely because it has been available as an R package and is easy to use [36-38]. The algorithmic modeling approach uses a distribution-free algorithm; examples include random forest classification, support vector machines, and artificial neural networks. The statistical community has developed and evaluated data modeling approaches [42]. In contrast, distribution-free algorithmic modeling, both in theory and practice, has developed rapidly in computational fields outside statistics. Algorithmic models are typically applied to large complex data sets and more recently have become known by plant breeders as machine learning algorithms.

Most comparative studies of GP methods have used continuous traits [40, 43-48]. While yield is clearly the most important trait for commodity crops such as maize, soybean and wheat, the majority of economically important traits, such as IDC, are evaluated on ordinal scales. Only a few comparative GP studies have used ordinal categorical phenotypes. These studies include GP of ordinal data using Bayesian logistic regression for maize gray leaf spot [49] and GP using several Bayesian models for resistance to pear black spot resistance [50]. Further, as far as we know, comparisons among GP models use *r* as a metric. However, plant breeders need more informative metrics. Plant breeding is a decision making discipline and decisions about germplasm are binary: retain or discard. Thus, in addition to *r*, decision metrics such as sensitivity, specificity, precision, decision accuracy and the area under the reciever operating characteric curve (defined in the Methods section below) should be reported in comparative GP studies.

Herein, we report a comparison of eight models using six metrics of relevance for making decisions about retention of lines in varietal development and genetic improvement projects. Two the methods, ridge regression, and ridge logistic regression represent data modeling approaches. Six of the methods, Naïve Bayes (NB), Random Forest (RF), K-nearest Neighbour (KNN), Support Vector Machine (SVM), Gradient Boost Machine (GBM), and Artificial Neural Network (ANN) are representatives of algorithmic modeling approaches.

## Materials and Methods

A total of 1,000 F_6:10_ from four years of a soybean breeding program were used for this study. These lines were in field trials from 2013 to 2016, with two replicates per location, and two to four locations per year. Phenotypic scores for IDC ranged from 1 to 9, with 1 as most resistant and 9 as most susceptible. The 1,000 lines were genotyped using a 1.5K Illumina chip of which 1,200 SNPs were of sufficient quality for use in the study. Missing genotypic data were imputed with the “impute” R package.

### Analytic methods

The best linear unbiased predicted (BLUP) value of IDC scores were estimated from multiple locations and years of data via the R package “lme4”. The mixed model for estimating the BLUP values for IDC: *IDC*_*score*_= µ + Year + Location + Line + experiment + Location x Line + Year x Line + e. All terms are treated as random effects except µ. BLUP IDC scores from the full data set were considered the true values against which GEBV’s in validation data sets created by a ten-fold cross validation approach for each of the GP methods.

Artificial neural network (ANN) used two hidden layers with the backdrop algorithm for predicting IDC scores, using the neuralnet package [51]. XGBoost is an R package [55] that is an optimized distributed gradient boosting machine (GBM) and it is an algorithm that has recently been dominating applied machine learning and Kaggle competitions for structured or tabular data[55, 56]. ridge regression, logistic ridge regression, Random forest (RF),, KNN, SVM, and Naïve Bayes (NB) analyses were conducted with the “rrBLUP”, “RDRR”, “randomForest”, “stats”, “class”, “e1071” and “naivebayes” R packages, respectively.

Prediction accuracy is estimated using the Pearson correlation coefficient determined as :

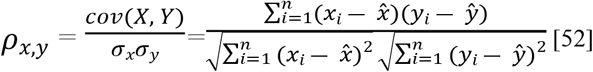

Where *cov*(*X, Y*) is covariance of X and Y, *σ*_*x*_ is the standard deviation of X, *σ*_*y*_ is the standard deviation for Y. 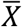 and 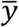 are the estimated means of X and Y, respectively. *x*_*i*_ *and y*_*i*_ are the observed values for X and Y.

In the context of decision making based on IDC scores, sensitivity is the estimated frequency of correctly identified tolerant scores [53]. The closer the sensitivity is to 1, the less risk of discarding tolerant lines. The estimated sensitivity is calculated as:

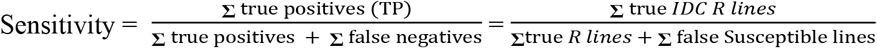

In the context of decision making based on IDC scores, specificity is the estimated frequency of correctly identified susceptible lines [53]. The closer the estimated specificity is to 1, the lower the risk of keeping susceptible lines in the varietal development and genetic improvement projects. Specificity is calculated as:

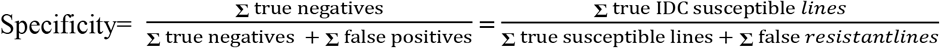

In the context of decision making based on IDC scores, precision is used to evaluate the ability to predict the resistant IDC lines with the risk of selecting the susceptible lines as resistant lines, reslting in wasted resources in the subsequent field trials and crossing nurseries. The higher the selection precision (closer to 1), the less risk of keeping susceptible lines in breeding program.

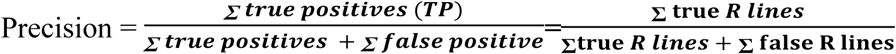

Decision accuracy

Decision accuracy is the proportion of true positives and true negative among all lines.

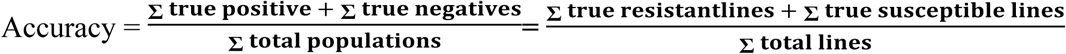

The receiver operating characteristic (ROC) curve is generated by plotting the true positive rate (TPR) on the y-axis against the false positive rate (FPR) on the x-axis at various decision threshold settings. The ROC curve is plotted with all values between 0 to 1. The Area under the ROC curve (AUC) is used to estimate the model’s ability to correctly inform decisions about retention and culling of lines. AUC values that are closer to 1 indicate better the models for making decisions [54, 55]. The AUC was calculated with the “ROCR” R package [56].

All R codes are available in a R markdown file in supplemental materials associated with this manuscript.

## Results and discussion

Averaged estimates of *r* from ten replicates of 10-fold cross validation data sets indicate that RR is better than LRR and three of the algorithmic models, but not as good as three of the algorithmic models (Table 1).

**Table 1.** Averaged estimates six metrics used to compare ten replicates of 10-fold cross validation data sets created by eight genomic prediction methods.

| GP method | AUC | Prediction Accuracy | Sensitivity | Specificity | Precision | Decision accuracy |
| --- | --- | --- | --- | --- | --- | --- |
| RR | 0.76 | 0.8739 | 0.9246 | 0.7510 | 0.7772 | 0.8528 |
| LRR | 0.51 | 0.5572 | 0.5112 | 0.5017 | 0.5088 | 0.5028 |
| SVM | 0.87 | 0.7233 | 0.6075 | 0.9665 | 0.9882 | 0.6710 |
| NB | 0.90 | 0.8448 | 0.9357 | 0.7405 | 0.7825 | 0.8379 |
| KNN | 0.47 | 0.8022 | 0.7874 | 0.8349 | 0.8280 | 0.8110 |
| ANN | 0.93 | 0.9110 | 0.9094 | 0.8981 | 0.8952 | 0.9020 |
| GBM | 0.95 | 0.9272 | 0.992 | 0.8904 | 0.8944 | 0.9090 |
| RF | 0.98 | 0.9491 | 0.9820 | 0.9340 | 0.9369 | 0.9580 |

Decision accuracy of SVM from replicated 10-fold cross-validation was 0.6710, which is lower than rrBLUP. However, it’s AUC, specificity and precision are better than RR. Selection sensitivity was 0.6075, which is the accuracy that true IDC resistance lines can be identified without false susceptible lines. The estimated sensitivity also indicates that about 40% of the resistant lines would be incorrectly discarded. In contrast the estimated specificity of 0.9665 and (Fig 1), indicates that we can safely discard the predicted susceptible IDC lines without discarding tolerant lines. High specificity means that the ability to identify the true susceptible IDC lines is much higher than that of identifying true resistant lines. High selection precision of 0.9882 indicated that the false-positives chance is very low and the true positive IDC resistance rate is high. The overall accuracy results from both specificity and precision showed that SVM can correctly identify susceptible lines with a much higher rate than identification of resistant lines. The SVM is very useful to exclude IDC susceptible lines during the early breeding stage rather than from subsequent resource-extensive yield testing.

The average decision accuracy for the NB GP model was 0.8370 (Table 1), which is similar to that from RR, 0.8528 (Table 1). The difference in decision accuracy values between NB and RR was not significant (p-value=0.06). But, in contrast to the results from SVM, which has very high discard specificity but low selection sensitivity rates, the NB model had a low specificity of 0.7405 but high sensitivity of 0.9357 (Fig 1). The difference level between the esimtated specificity and sensitivity values for NB are significant with p-value = 8.519e-07. These results indicate that NB can correctly predict the tolerant IDC lines better than it can predict susceptible lines. For some breeding situations it may be appropriate to conduct both SVM and NB genomic predictions, to correctly identify both resistant and susceptible IDC lines.

The averaged decision accuracy from10-fold cross-validation sets created by the KNN model is 0.8110 which is significantly (p = 5.137e-05) less than that of RR (Table 1 and Fig 1). The difference between estimates of sensitivity and specificity of the KNN model was not significant with p = 0.08058.

The averaged sensitivity, specificity, precision, and decision accuracies by applying ANN to validation sets created by replicated 10-fold cross-validation are 0.9094, 0.8981, 0.8952, and 0.9020, respectively (Table 1). There were no significant differences among these four metrics (p-values > 0.05). But, the average decision accuracy from the ANN GP model is significantly greater than RR (p<0.01; Fig 1). Upon closer inspection we note that the estimated standard deviations among replicated ten-fold cross validations using ANN are consistently small values for all decision metrics indicating that the model provides stable results overfitting is unlikely.

Results from GBM were similar to those from ANN. The average sensitivity, specificity, selection precision, and decision accuracies created by XGBoost-GBM from the 10-fold cross-validation data sets were consistently close to 0.90 (Table 71). The difference among these four metrics were not significant with p-values > 0.05. The difference t-test was significant with a p-value of 0.0118 (<0.05) when comparing ANN with RR (Fig 1). Even though these two means from 10-fold cross-validation, 0.909 and 0.8739, are very close, their difference is significant.

Results from application of the RF GP method to IDC scores were the best of all GP methods. The averaged estimates of sensitivity, specificity, precision, and decision accuracies from 10-fold cross-validations were 0.9340, 0.9820, 0.9369, and 0.9580, respectively (Table 1). The average specificity was significantly larger than the sensitivity (p = 9.88e-05). The p-value of the t-test between the decision accuracies generated by RF and rrBLUP was 1.344e-07.

The combined results of the six algorithmic models can be summarized as follows. The best prediction accuracy was obtained using the SVM (0.9886). The SVM also provided the best ability to correctly discard lines that are susceptible to IDC (0.9713). Thus, the SVM provides the best information for the breeder to avoid retaining lines and wasting resources in subsequent phases of varietal development and recurrent genetic improvement projects. The RF method provided the best ability to identify tolerant lines (0.982) for retention in varietal development and genetic improvement projects. Decision accuracies from ANN, GBM, and RF were all higher than that from RR. ANN and RF have the smallest boxplot height between the Q1 25% and Q3 75% quantile, which indicates performance of ANN and RF are very stable across cross-validations (Fig 2). In contrast, the height of the boxplot from SVM is the largest, which shows that the accuracy from SVM varies the most and not stable. Penalized Logistic regression provided the least accurate decisions. Decision accuracies of ANN, GBM, and RF, 0.902, 0.909, 0.958, respectively, are significantly greater than RR (0.8739). For IDC scores provided by four years of IDC field trials conducted by a commercial organization algorithmic models outperform the data models.

**Fig 2.**
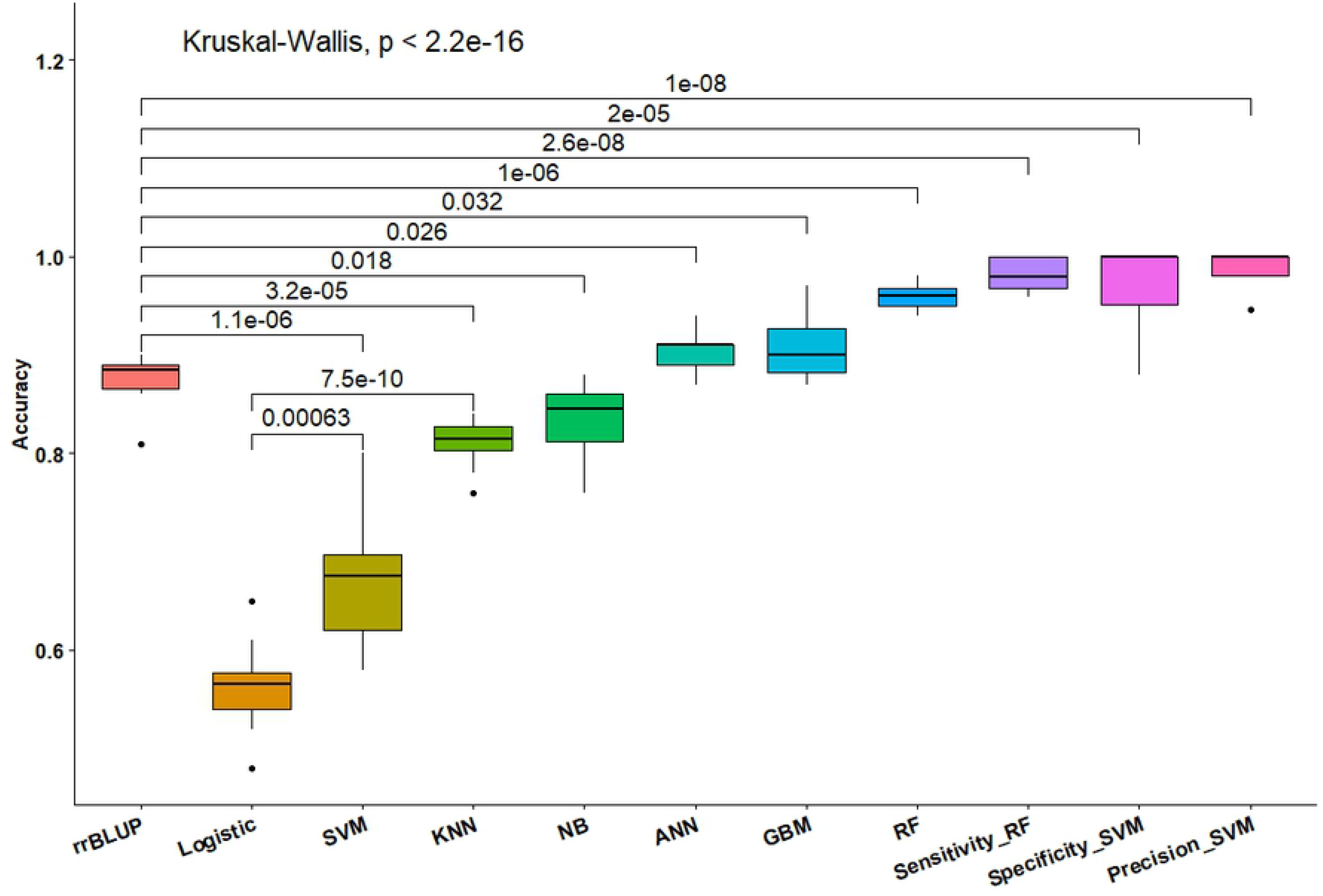
Comparisons among eight models. Y-axis are the coefficients of efficiency. Black dots are the outliers, and the numbers above each line are the p-values of the significant pairwise tests. The value at the top of this Fig is the overall p-value from the Kruskal-Wallis test.

Results from the AUC values (Table 1) and ROC curves from all the data of each model show that RF is the most stable model with the lowest standard deviation of 0.014. ANN, GBM, and RF are very similar in regard to both ROC and estimated standard deviations. In contrast, KNN and LRR have small ROC values, indicating both models face challenges of balancing false positive and false negative scores when attempting to distinguish tolerant from susceptible IDC lines (Fig 3). The AUC of SVM is lower than that of RF, which is consistent with overall low accuracy but higher specificity from SVM. KNN is the poorest model with the lowest AUC value and highest standard deviation.

**Fig 3.**
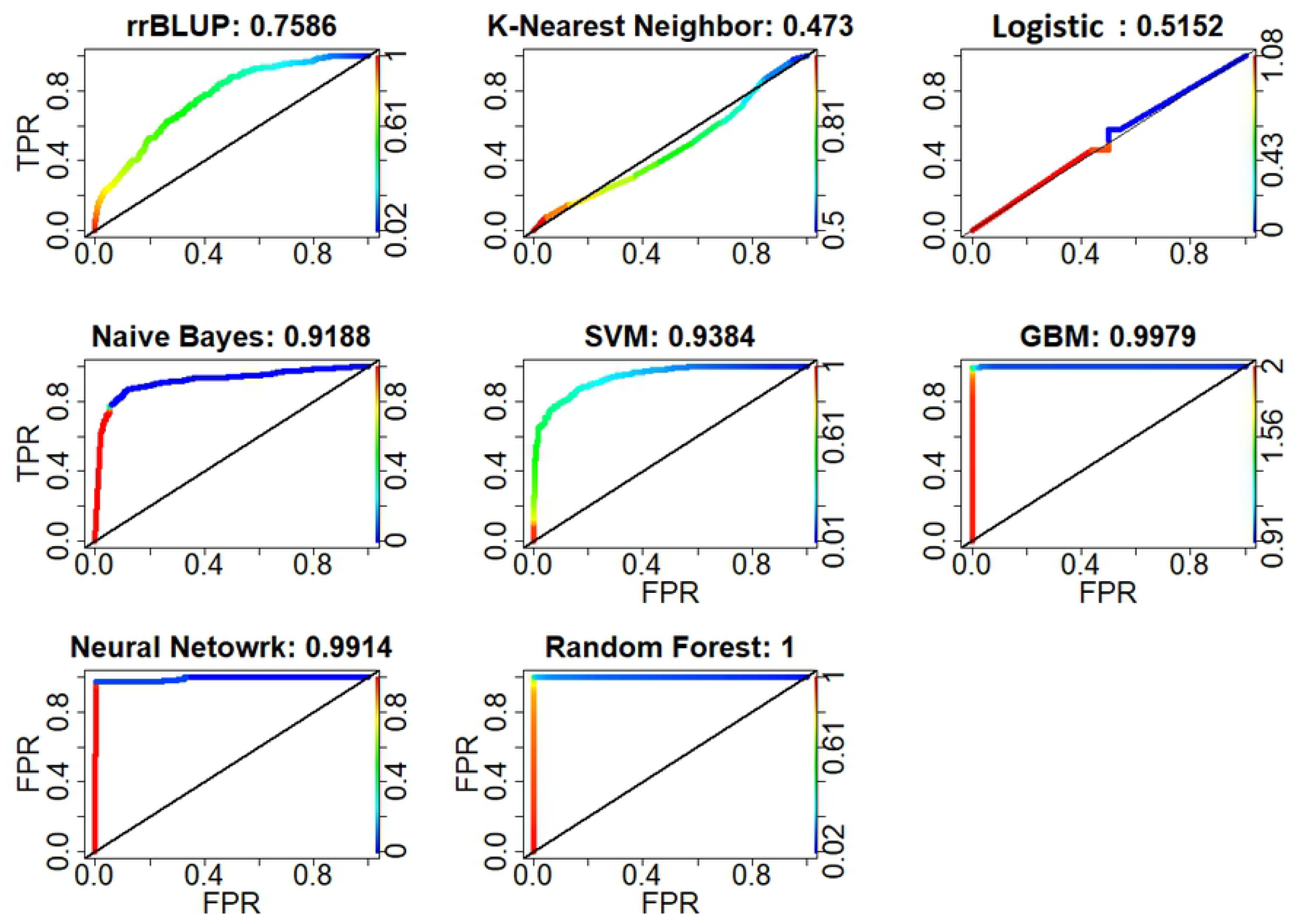
Area under curve (AUC) of the receiver operating characteristic (ROC) curve from eight models. X-axis is false positive rate (FPR), left y-axis is true positive rate (TPR), right y-axis are cutoffs used to calculate the false and true positive rates with green for small and red for large cutoff values. On the top of each image the number after the colon is the AUC value, from 0 to 1.

The 45-degree diagonal line of each AUC is the baseline from random classifications with TPR=FPR. The color of the curve is a mixture of green for TPR and red for FPR.

Results from the significant pairwise t-test of the AUC values show that LRR and KNN provide similar decision accuracies (p = 0.088; Fig 4) and they are the worst among the eight models. The SVM, NB, ANN, GBM, and RF have significantly greater (p-values < 0.001) AUC values than RR. Overall, the algorithmic models outperform the data models in regard to the AUCs.

**Fig 4.**
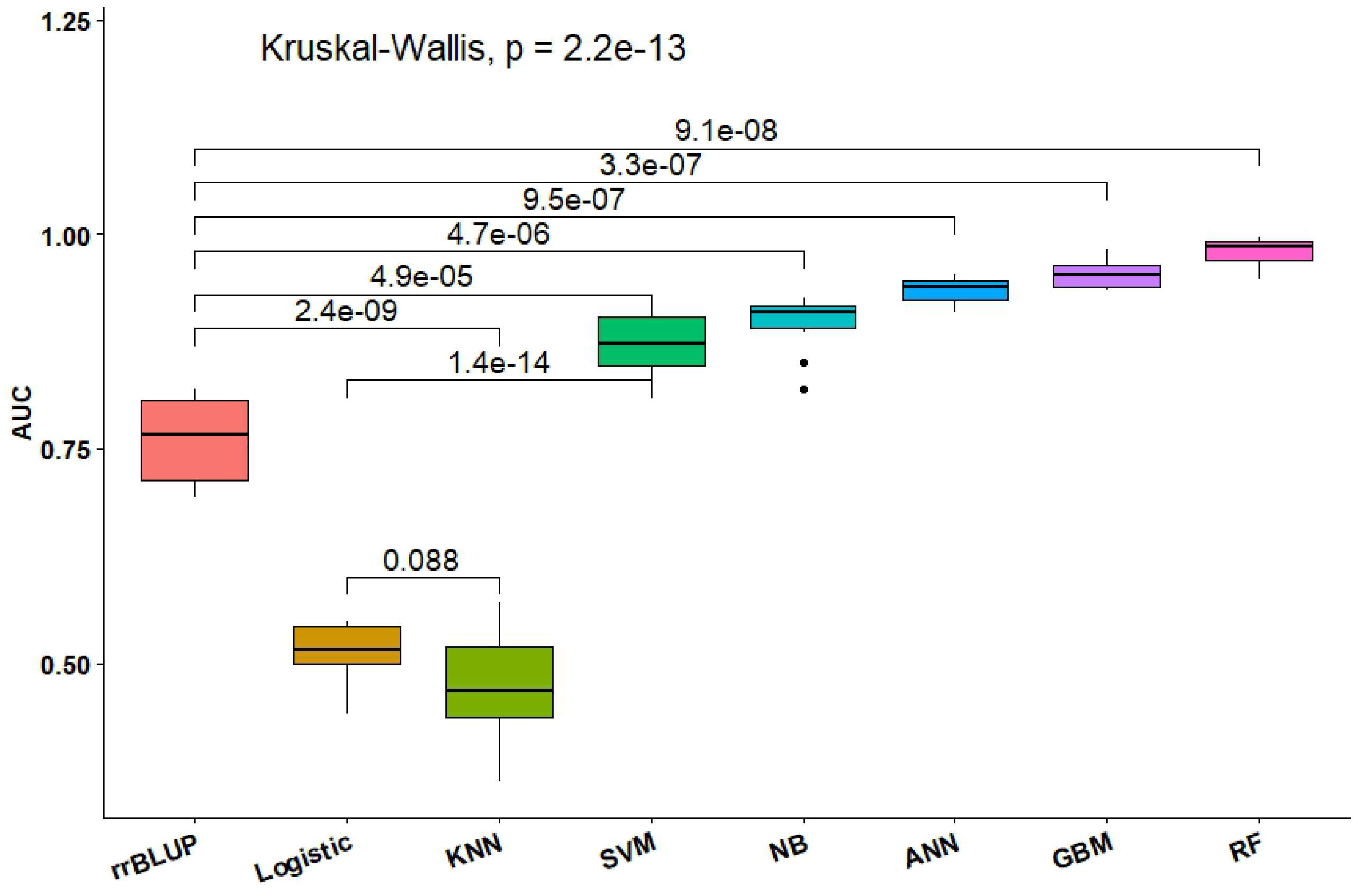
Summary of pairwise significant and overall Kruskal-Wallis test results. The values above horizontal lines are the p-values of the significant pairwise tests. The horizontal lines inside each box represent the means. The black dots are the outliers.

Results from a PCA analysis using IDC sensitivity, specificity, precision, decision accuracy, and AUC (Fig 5) cluster LRR and RR, while NB is separated from ANN, GBM, and RFcluster together.

**Fig 5.**
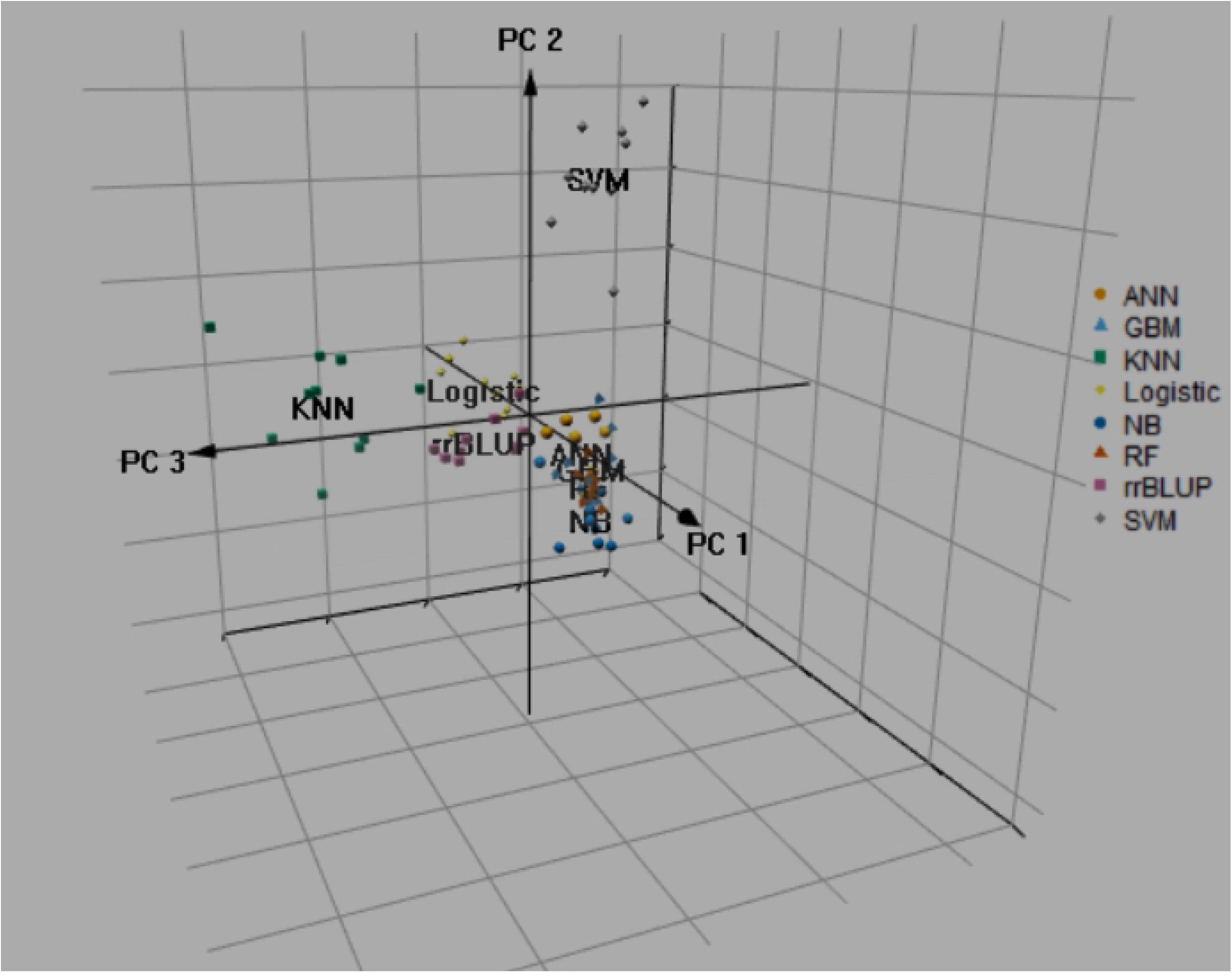
Display of the eight models by Principal Component Analysis (PCA). Features used for the PCA analysis were the IDC selection sensitivity, discard specificity, selection precision, prediction overall accuracy, and AUC values.

Results from a non-parametric Kruskal-Wallis test of estimated averages of the AUC and decision accuracy values from each of the models showed that means of objective-specific selection sensitivity with RF, discard specificity with SVM, and selection precision with SVM for IDC were significantly better than other models (Table 2). From these observations, different models should be applied for selecting resistant and culling susceptible lines in breeding for IDC. RF is the best model with combined accuracy and AUC values. ANN and GBM are the third best models. In contrast, RR is about the same as SVM. The LRR ranked last in the multiple comparisons and group analysis. Compared with RR, precision using SVM and specificity using RF are better GP models to use for IDC.

**Table 2.** Results of Kruskal-Wallis test and mean separation from AUC and accuracy.

| Models | Overall mean * | Kruskal-Wallis ranking | Groups ($\alpha=0.05$) |
| --- | --- | --- | --- |
| RF | 0.969 | 154.9 | b |
| GBM | 0.931 | 125.1 | c |
| ANN | 0.918 | 114.2 | c |
| NB | 0.866 | 84.5 | d |
| RR | 0.819 | 66.4 | e |
| SVM | 0.771 | 61.2 | e |
| KNN | 0.641 | 33.8 | f |
| LRR | 0.537 | 18.7 | g |

\*: Overall mean of the 10 AUC and 10 accuracy values from each of the models.

## Conclusions

Among the eight tested models, the SVM provided the best ability to discard IDC susceptible lines, while the RF method was best for accuratly selecting IDC resistant lines with the greatest AUC. In an overall comparison of GP models applied to ordinal soybean IDC scores, the algorithmic modeling approaches outperformed the data modeling approaches, consistent with Breiman’s conclusions about the two statistical modeling cultures [42].

## Supplemental Data Available

Supplemental material is available online for this article. R codes and the data sets used for this research are available online at http://gfspopgen.agron.iastate.edu/resources.html

## Acknowledgments

We thank Iowa State University for the financial support of this research and thank Dr. Deborah Samac for her helpful comments on the manuscript. This research was supported in part by the US. Department of Agriculture, Agricultural Research Service, project 5030-21000-062-00D. Mention of trade names or commercial products in this publication is solely for the purpose of providing specific information and does not imply recommendation or endorsement by the U.S. Department of Agriculture. USDA is an equal opportunity provider and Employer.

## References

1. Hansen NC, Jolley VD, Naeve SL, Goos RJ. Iron deficiency of soybean in the North Central U.S. and associated soil properties. Iron deficiency of soybean in the North Central US and associated soil properties. 2004;(7):983–7.

2. Cianzio SRd, Fehr WR. Genetic control of iron deficiency chlorosis in soybeans. Iowa State Journal of Research. 1980;54(3):367–75. PubMed PMID: CABI:19811695504.

3. Lin S, Cianzio S, Shoemaker R. Mapping genetic loci for iron deficiency chlorosis in soybean. Molecular Breeding. 1997;3(3):219–29. doi: 10.1023/a:1009637320805. PubMed PMID: WOS:A1997XB24500008.

4. Lin SF, Grant D, Cianzio S, Shoemaker R. Molecular characterization of iron deficiency chlorosis in soybean. Journal of Plant Nutrition. 2000;23(11-12):1929–39. doi: 10.1080/01904160009382154. PubMed PMID: WOS:000166500800033.

5. Charlson DV, Cianzio SR, Shoemaker RC. Associating SSR markers with soybean resistance to iron deficiency chlorosis. Journal of Plant Nutrition. 2003;26(10-11):2267–76. doi: 10.1081/pln-120024280. PubMed PMID: WOS:000185604600030.

6. O’Rourke JA, Charlson DV, Gonzalez DO, Vodkin LO, Graham MA, Cianzio SR, et al. Microarray analysis of iron deficiency chlorosis in near-isogenic soybean lines. Bmc Genomics. 2007;8. doi: 10.1186/1471-2164-8-476. PubMed PMID: WOS:000253467400002.

7. Wang J, McClean PE, Lee R, Goos RJ, Helms T. Association mapping of iron deficiency chlorosis loci in soybean (Glycine max L. Merr.) advanced breeding lines. Theoretical and Applied Genetics. 2008;116(6):777–87. doi: 10.1007/s00122-008-0710-x. PubMed PMID: WOS:000254262600004.

8. Mamidi S, Chikara S, Goos RJ, Hyten DL, Annam D, Moghaddam SM, et al. Genome-Wide Association Analysis Identifies Candidate Genes Associated with Iron Deficiency Chlorosis in Soybean. Plant Genome. 2011;4(3):154–64. doi: 10.3835/plantgenome2011.04.0011. PubMed PMID: WOS:000312661700001.

9. King KE, Peiffer GA, Reddy M, Lauter N, Lin SF, Cianzio S, et al. Mapping of iron and zinc quantitative trait loci in soybean for association to iron deficiency chlorosis resistance. Journal of Plant Nutrition. 2013;36(14):2132–53. doi: 10.1080/01904167.2013.766804. PubMed PMID: WOS:000326070100002.

10. Xu Z, Cannon SB, Beavis WD. Applications of spatial models to ordinal data. bioRxiv. 2020:2020.09.21.306001. doi: 10.1101/2020.09.21.306001.

11. Mitchum MG. Soybean Resistance to the Soybean Cyst Nematode Heterodera glycines: An Update. Phytopathology. 2016;106(12):1444. doi: 10.1094/PHYTO-06-16-0227-RVW.

12. Fehr WR. Control of iron-deficiency chlorosis in soybeans by plant-breeding. Journal of Plant Nutrition. 1982;5(4-7):611–21. doi: 10.1080/01904168209362989. PubMed PMID: WOS:A1982NX30700037.

13. Cianzio SR, Fehr WR. Variation in the inheritance of resistance to iron-deficiency chlorosis in soybeans. Crop Science. 1982;22(2):433–4. PubMed PMID: WOS:A1982NQ29700051.

14. O’Rourke JA, Nelson RT, Grant D, Schmutz J, Grimwood J, Cannon S, et al. Integrating microarray analysis and the soybean genome to understand the soybeans iron deficiency response. BMC Genomics. 2009;10:376. doi: 10.1186/1471-2164-10-376. PubMed PMID: 19678937; PubMed Central PMCID: PMCPMC2907705.

15. Jannink JL, Lorenz AJ, Iwata H. Genomic selection in plant breeding: from theory to practice. Brief Funct Genomics. 2010;9(2):166–77. doi: 10.1093/bfgp/elq001. PubMed PMID: 20156985.

16. Rodriguez de Cianzio S, Fehr WR. Variation in the inheritance of resistance to iron deficiency chlorosis in soybeans. Crop Science. 1982;22(2):433–4. PubMed PMID: CABI:19821615691.

17. Meuwissen TH, Hayes BJ, Goddard ME. Prediction of total genetic value using genome-wide dense marker maps. Genetics. 2001;157(4):1819–29. PubMed PMID: 11290733; PubMed Central PMCID: PMCPMC1461589.

18. Bernardo R. Molecular markers and selection for complex traits in plants: Learning from the last 20 years. Crop Science. 2008;48(5):1649–64. doi: 10.2135/cropsci2008.03.0131. PubMed PMID: WOS:000259792100001.

19. Heffner EL, Jannink J-L, Iwata H, Souza E, Sorrells ME. Genomic Selection Accuracy for Grain Quality Traits in Biparental Wheat Populations. Crop Science. 2011;51(6):2597–606. doi: 10.2135/cropsci2011.05.0253. PubMed PMID: WOS:000295839200031.

20. Heffner EL, Jannink J-L, Sorrells ME. Genomic Selection Accuracy using Multifamily Prediction Models in a Wheat Breeding Program. Plant Genome. 2011;4(1):65–75. doi: 10.3835/plantgenome2010.12.0029. PubMed PMID: WOS:000312661400007.

21. Heffner EL, Lorenz AJ, Jannink J-L, Sorrells ME. Plant Breeding with Genomic Selection: Gain per Unit Time and Cost. Crop Science. 2010;50(5):1681–90. doi: 10.2135/cropsci2009.11.0662. PubMed PMID: WOS:000281060300010.

22. Gorjanc G, Jenko J, Hearne SJ, Hickey JM. Initiating maize pre-breeding programs using genomic selection to harness polygenic variation from landrace populations. Bmc Genomics. 2016;17. doi: 10.1186/s12864-015-2345-z. PubMed PMID: WOS:000367566300013.

23. Lian L, Jacobson A, Zhong S, Bernardo R. Prediction of Genetic Variance in Biparental Maize Populations: Genomewide Marker Effects versus Mean Genetic Variance in Prior Populations. Crop Science. 2015;55(3):1181–8. doi: 10.2135/cropsci2014.100729. PubMed PMID: WOS:000357779400021.

24. Iwata H, Jannink J-L. Accuracy of Genomic Selection Prediction in Barley Breeding Programs: A Simulation Study Based On the Real Single Nucleotide Polymorphism Data of Barley Breeding Lines. Crop Science. 2011;51(5):1915–27. doi: 10.2135/cropsci2010.12.0732. PubMed PMID: WOS:000293473800004.

25. Sallam AH, Endelman JB, Jannink JL, Smith KP. Assessing Genomic Selection Prediction Accuracy in a Dynamic Barley Breeding Population. Plant Genome. 2015;8(1). doi: 10.3835/plantgenome2014.05.0020. PubMed PMID: WOS:000352494900001.

26. Schmid KJ, Thorwarth P. Genomic Selection in Barley Breeding. In: Kumlehn J, Stein N, editors. Biotechnological Approaches to Barley Improvement. Biotechnology in Agriculture and Forestry. 692014. p. 367–78.

27. Muranty H, Troggio M, Ben Sadok I, Al Rifai M, Auwerkerken A, Banchi E, et al. Accuracy and responses of genomic selection on key traits in apple breeding. Horticulture Research. 2015;2. doi: 10.1038/hortres.2015.60. PubMed PMID: WOS:000367188400001.

28. Hayes BJ, Bowman PJ, Chamberlain AJ, Goddard ME. Invited review: Genomic selection in dairy cattle: progress and challenges (vol 92, pg 433, 2009). Journal of Dairy Science. 2009;92(3):1313-. doi: 10.3168/jds.2009-92-3-1313. PubMed PMID: WOS:000263559600051.

29. Montaldo HH, Casas E, Ferraz JBS, Vega-Murillo VE, Roman-Ponce SI. Opportunities and challenges from the use of genomic selection for beef cattle breeding in Latin America. Animal Frontiers. 2012;2(1):23–9. doi: 10.2527/af.2011-0029. PubMed PMID: CABI:20123393808.

30. Hayes BJ, Daetwyler HD, Bowman P, Moser G, Tier B, Crump R, et al. Accuracy of genomic selection: comparing theory and results2009. 34–7 p.

31. Daetwyler HD, Villanueva B, Woolliams JA. Accuracy of Predicting the Genetic Risk of Disease Using a Genome-Wide Approach. Plos One. 2008;3(10). doi: 10.1371/journal.pone.0003395. PubMed PMID: WOS:000265121700004.

32. Lian L, Jacobson A, Zhong S, Bernardo R. Genomewide Prediction Accuracy within 969 Maize Biparental Populations. Crop Science. 2014;54(4):1514–22. doi: 10.2135/cropsci2013.12.0856. PubMed PMID: WOS:000338773100022.

33. Lorenz AJ. Resource Allocation for Maximizing Prediction Accuracy and Genetic Gain of Genomic Selection in Plant Breeding: A Simulation Experiment. G3-Genes Genomes Genetics. 2013;3(3):481–91. doi: 10.1534/g3.112.004911. PubMed PMID: WOS:000315950000009.

34. Lorenz AJ, Smith KP. Adding Genetically Distant Individuals to Training Populations Reduces Genomic Prediction Accuracy in Barley. Crop Science. 2015;55(6):2657–67. doi: 10.2135/cropsci2014.12.0827. PubMed PMID: WOS:000368265600025.

35. Sallam AH, Endelman JB, Jannink JL, Smith KP. Assessing genomic selection prediction accuracy in a dynamic barley breeding population. Plant Genome. 2015;8(1):unpaginated-unpaginated. PubMed PMID: CABI:20153126029.

36. Endelman JB. Ridge Regression and Other Kernels for Genomic Selection with R Package rrBLUP. Plant Genome. 2011;4(3):250–5. doi: 10.3835/plantgenome2011.08.0024. PubMed PMID: WOS:000312661700009.

37. Nishio M, Satoh M. Including dominance effects in the genomic BLUP method for genomic evaluation. PLoS One. 2014;9(1):e85792. doi: 10.1371/journal.pone.0085792. PubMed PMID: 24416447; PubMed Central PMCID: PMCPMC3885721.

38. Perez-Rodriguez P, Gianola D, Weigel KA, Rosa GJ, Crossa J. Technical note: An R package for fitting Bayesian regularized neural networks with applications in animal breeding. J Anim Sci. 2013;91(8):3522–31. doi: 10.2527/jas.2012-6162. PubMed PMID: 23658327.

39. Covarrubias-Pazaran G. Genome-Assisted Prediction of Quantitative Traits Using the R Package sommer. PLoS One. 2016;11(6):e0156744. doi: 10.1371/journal.pone.0156744. PubMed PMID: 27271781; PubMed Central PMCID: PMCPMC4894563.

40. Lorenzana RE, Bernardo R. Accuracy of genotypic value predictions for marker-based selection in biparental plant populations. Theoretical and Applied Genetics. 2009;120(1):151–61. doi: 10.1007/s00122-009-1166-3. PubMed PMID: WOS:000271939900013.

41. Cros D, Denis M, Sanchez L, Cochard B, Flori A, Durand-Gasselin T, et al. Genomic selection prediction accuracy in a perennial crop: case study of oil palm (Elaeis guineensis Jacq.). Theoretical and Applied Genetics. 2015;128(3):397–410. doi: 10.1007/s00122-014-2439-z. PubMed PMID: WOS:000350041500003.

42. Breiman L. Statistical modeling: The two cultures. Statistical Science. 2001;16(3):199–215. doi: 10.1214/ss/1009213726. PubMed PMID: WOS:000172846900001.

43. Ogutu JO, Piepho H-P, Schulz-Streeck T. A comparison of random forests, boosting and support vector machines for genomic selection. BMC proceedings. 2011;5 Suppl 3:S11–S. doi: 10.1186/1753-6561-5-s3-s11. PubMed PMID: MEDLINE:21624167.

44. Ogutu JO, Schulz-Streeck T, Piepho HP. Genomic selection using regularized linear regression models: ridge regression, lasso, elastic net and their extensions. BMC Proc. 2012;6 Suppl 2:S10. doi: 10.1186/1753-6561-6-S2-S10. PubMed PMID: 22640436; PubMed Central PMCID: PMCPMC3363152.

45. Shu Y, Wu L, Wang D, Guo C. Application of artificial neural network in genomic selection for crop improvement. Acta Agronomica Sinica. 2011;37(12):2179–86. PubMed PMID: CABI:20123022436.

46. Hayes BJ, Bowman PJ, Chamberlain AJ, Goddard ME. Invited review: Genomic selection in dairy cattle: Progress and challenges. Journal of Dairy Science. 2009;92(2):433–43. doi: 10.3168/jds.2008-1646. PubMed PMID: WOS:000262654900001.

47. Ratcliffe B, El-Dien OG, Klapste J, Porth I, Chen C, Jaquish B, et al. A comparison of genomic selection models across time in interior spruce (Picea engelmannii x glauca) using unordered SNP imputation methods. Heredity. 2015;115(6):547–55. doi: 10.1038/hdy.2015.57. PubMed PMID: WOS:000365123300009.

48. Zargar SM, Raatz B, Sonah H, Muslima N, Bhat JA, Dar ZA, et al. Recent advances in molecular marker techniques: insight into QTL mapping, GWAS and genomic selection in plants. Journal of Crop Science and Biotechnology. 2015;18(5):293–308. doi: 10.1007/s12892-015-0037-5. PubMed PMID: CABI:20163024709.

49. Montesinos-Lopez OA, Montesinos-Lopez A, Crossa J, Burgueno J, Eskridge K. Genomic-Enabled Prediction of Ordinal Data with Bayesian Logistic Ordinal Regression. G3-Genes Genomes Genetics. 2015;5(10):2113–26. doi: 10.1534/g3.115.021154. PubMed PMID: WOS:000362505600019.

50. Iwata H, Hayashi T, Terakami S, Takada N, Sawamura Y, Yamamoto T. Potential assessment of genome-wide association study and genomic selection in Japanese pear Pyrus pyrifolia. Breeding Science. 2013;63(1):125–40. doi: 10.1270/jsbbs.63.125. PubMed PMID: WOS:000322417000015.

51. Günther F, Fritsch S. neuralnet: Training of Neural Networks. The R Journal. 2010;2(1):30. doi: 10.32614/RJ-2010-006.

52. Ott L, Longnecker M. An introduction to statistical methods and data analysis. 6th ed. Belmont, CA: Brooks/Cole Cengage Learning; 2010. xv, 1273 p. p.

53. Gaddis G, Gaddis M. Introduction to Biostatistics: Part 3,Sensitivity, Specificity, Predictive Value,and Hypothesis Testing. Annals of emergency medicine. Lansing, Mich.: American College of Emergency Physicians.; 1990. p. 145–51.

54. Pepe MS, Cai T, Longton G. Combining Predictors for Classification Using the Area under the Receiver Operating Characteristic Curve. Combining Predictors for Classification Using the Area under the Receiver Operating Characteristic Curve. 2006;62(1):221–9. doi: 10.1111/j.1541-0420.2005.00420.x.

55. Peterson LE, Coleman MA. Machine learning-based receiver operating characteristic (ROC) curves for crisp and fuzzy classification of DNA microarrays in cancer research. International Journal of Approximate Reasoning. 2008;47(1):17–36. doi: 10.1016/j.ijar.2007.03.006.

56. Sing T, Sander O, Beerenwinkel N, Lengauer T. ROCR: visualizing classifier performance in R. Bioinformatics. 2005;21(20):3940–1. doi: 10.1093/bioinformatics/bti623.

57. Jeffrey BE. Ridge Regression and Other Kernels for Genomic Selection with R Package rrBLUP. The Plant Genome. 2011;4(3):250–5. doi: 10.3835/plantgenome2011.08.0024.

58. Ogutu JO, Schulz-Streeck T, Piepho H-P. Genomic selection using regularized linear regression models: ridge regression, lasso, elastic net and their extensions. BMC proceedings. 2012;6 Suppl 2(Suppl 2):S10. doi: 10.1186/1753-6561-6-S2-S10.

59. Piepho HP, Ogutu JO, Schulz-Streeck T, Estaghvirou B, Gordillo A, Technow F. Efficient Computation of Ridge-Regression Best Linear Unbiased Prediction in Genomic Selection in Plant Breeding. Efficient Computation of Ridge-Regression Best Linear Unbiased Prediction in Genomic Selection in Plant Breeding. 2012;52(3):1093–104. doi: 10.2135/cropsci2011.11.0592.

